# Cortical F-actin filament organization corrals E-cadherin/HMR-1 during cytokinetic furrow ingression in *C. elegans* embryos

**DOI:** 10.1101/2023.12.07.570732

**Authors:** Debodyuti Mondal, Megha Rai, Anup Padmanabhan

## Abstract

In metazoan cells, the delicate equilibrium between proliferative cell division and inter-cellular adhesion is crucial for achieving growth while maintaining tissue integrity and homeostasis. The transmembrane adhesion receptor, E-cadherin, not only facilitates cell-cell adhesion but modulates cytokinesis by impeding cortical F-actin flow and inhibiting actomyosin contractility. In the *C. elegans* zygote, newly established cell-cell interfaces following cytokinesis lack E-cadherin/HMR-1 (Padmanabhan et al., 2017b). The mechanisms responsible for the spatial exclusion of HMR-1, however remain elusive. Here, we show that HMR-1 is actively prevented from entering the cytokinetic furrow zone during early embryonic divisions in *C. elegans*. The compact alignment of unbranched F-actin filaments polymerized by the formin/CYK-1 restricts HMR-1 from entering the ingressing furrow. This corralling phenomenon depends on the physical association between HMR-1 and F-actin mediated via the β- and α- catenins, HMP-2 and HMP-1, respectively. Our findings reveal a previously unknown reciprocal regulatory relationship between cell adhesion and cell division machineries, ensuring the precise orchestration of the cytokinesis during embryonic development.

## INTRODUCTION

Eukaryotic cytokinesis relies on the constriction of actomyosin based contractile ring that bisects the segregating genetic material. Studies carried out across various species have catalogued the core components that make up the cytokinetic ring, revealing conserved genes from yeast to metazoans(D’Avino et al., 2004). The assembly of the cytokinetic apparatus in animal cells is initiated upon anaphase onset, following the activation of the small GTPase RhoA. Localized RhoA activation initiates the recruitment and/or assembly of key components of the cytokinetic ring, including, type-II non-muscle myosin, diaphanous formin, anillin, septins, and acto-myosin binding and regulatory proteins (Basant and Glotzer, 2018; Mierzwa and Gerlich, 2014). The motor activity of myosin on F-actin filaments enables constriction of the cytokinetic apparatus that drives the ingression of the cell cortex.

While significant progress in our understanding of cytokinesis has been made through studies in isolated cells, in metazoans, intercellular adhesion has been implicated in the regulation of the plane and pace of cytokinetic furrow ingression (Derksen and van de Ven, 2020; Founounou et al., 2013; Guillot and Lecuit, 2013; Herszterg et al., 2013; Morais-De-Sá and Sunkel, 2013; Padmanabhan et al., 2017b; Pinheiro et al., 2017). The transmembrane adhesion receptor-E-cadherin, is essential for embryonic development and epithelial tissue integrity in metazoans (Bulgakova et al., 2012; Costa et al., 1998; Costa et al., 1998; Harris and Tepass, 2010; Ninomiya et al., 2012; Takeichi, 2014). E-cadherin mediated adhesion requires receptors expressed on the cell surface to interact in-trans through their extracellular domain. Cytoplasmic adaptor proteins – β-catenin and ⍺-catenin are recruited to the intracellular domain of E-cadherin. ⍺-catenin in-turn binds cortical F-actin via its actin binding domains. In addition to core components, a complex of ∼150 proteins are recruited to the sites of cell adhesion (Bertocchi et al., 2017; Guo et al., 2014). Intercellular adhesion is intricately linked with other physiological processes such as cellular proliferation, migration, or apoptosis. Therefore, spatiotemporal plasticity in remodeling of junctional E-cadherin in the adhesion machinery is necessary to ensure accuracy of cell divisions during proliferative tissue growth and morphogenesis (Takeichi, 2014; (Founounou et al., 2013; Guillot and Lecuit, 2013; Herszterg et al., 2013).

While E-cadherin is predominantly localized to cell junctions mediating inter-cellular adhesion, a significant population of E-cadherin is also found at non-junctional surfaces (Padmanabhan et al., 2017b; Wu et al., 2015). We have previously shown that during early embryonic divisions in *C. elegans* embryos, the E-cadherin ortholog HMR-1, in addition to associating with the F-actin, inhibits RHO-1 dependent cortical myosin contractility, slowing down cytokinetic furrow ingression. This indicated an inverse relationship between E-cadherin/HMR-1 and actomyosin dependent cytokinesis machinery. An observation that HMR-1 was never detected in the newly formed cell boundary following cytokinesis, prompted us to ask if there existed a reciprocal regulation where the contractile actomyosin ring regulates the surface localization and dynamics of HMR-1 during cell division.

Here we show that in *C. elegans* embryonic divisions, orthogonally aligned cortical F-actin filaments polymerized by formin/CYK-1 functions as fence bordering the contractile furrow zone. The cortical flows during cytokinetic furrow ingression causes surface localized HMR-1 clusters to associate with these F-actin filaments, via catenin adaptors - HMP-2 and HMP-1. This association between HMR-1 clusters and the cortical architecture of F-actin filaments in the furrow zone confines and limits the movement of HMR-1 into the furrow. Our findings provide evidence supporting a potential mechanism for reciprocal inhibition between E-cadherin mediated cell adhesion and actomyosin based cell division machinery.

## RESULTS

### Surface localized HMR-1 clusters are restricted from the medial furrow zone during cytokinesis

Following anaphase onset, the circumferentially assembled contractile actomyosin ring, constricts to divide the cell. The non-muscle type-II myosin (NMY-2) enriched at edge of the ingressing furrow, is the primary force generator driving cytokinesis (D’Avino et al., 2015; Green et al., 2012). To characterize the dynamics of HMR-1 as a new boundary forms between daughter cells during cytokinesis, we carried out confocal microscopy of the surface plane of dividing embryos co-expressing GFP fused to the C-terminus of HMR-1 (HMR-1::GFP) and the plasma membrane marker, mCherry fused to PLC-1ι-PH (mCherry::PH). As observed earlier, HMR-1 receptors aggregate at inter cellular junctions (Fig 1A; yellow arrowheads in AB and P1 division) as well as at non-junctional surfaces (Fig 1A; cyan arrowheads) of early embryonic cells in *C. elegans* (Padmanabhan et al., 2017b). However, post anaphase onset in P0, AB and P1 cells, HMR-1 clusters in were observed in the non-furrowing regions of the cellular surface but absent in the contractile furrow zone (Fig 1A; red region of interest ROI). The double membrane formed due to the plasma membrane folding in during furrow ingression of P0 cells lacked HMR-1 (Top panel in Fig 1B and S1A, yellow arrowheads) (Padmanabhan et al., 2017b). We then checked if the absence of HMR-1 in the furrow could be observed subsequent embryonic divisions. Like the P0 zygote, we did not detect HMR-1 in the ingressing furrows of AB and P1 divisions, (Fig S1B, yellow arrowhead). HMR-1 receptors are excluded from the furrow zone throughout cytokinesis and localizes to the newly formed cell interface only after the furrow ingression is completed (Fig S1A and S1B, yellow arrow). To check if the spatial inhibition at the furrow region is specific to HMR-1 or general phenomena among transmembrane proteins, we imaged the first and second divisions of embryos expressing mScarlet tagged PEZO-1 (mScar::PEZO-1), the multi transmembrane calcium gated mechanosensory protein and GFP tagged SAX-7 (SAX-7::eGFP), the L1-IgCAM cell adhesion receptor (Bai et al., 2020; Chen et al., 2001; Wang et al., 2005). In contrast to the HMR-1, both mScar::PEZO-1 and SAX-7::eGFP localized to furrows throughout ingression during P0 and AB divisions (Fig 1B, yellow arrows, (Chen et al., 2001)). Thus, the spatial exclusion from the furrow zone is specifically exhibited by HMR-1, and not by other cell adhesion receptors or transmembrane proteins.

**Figure 1:**
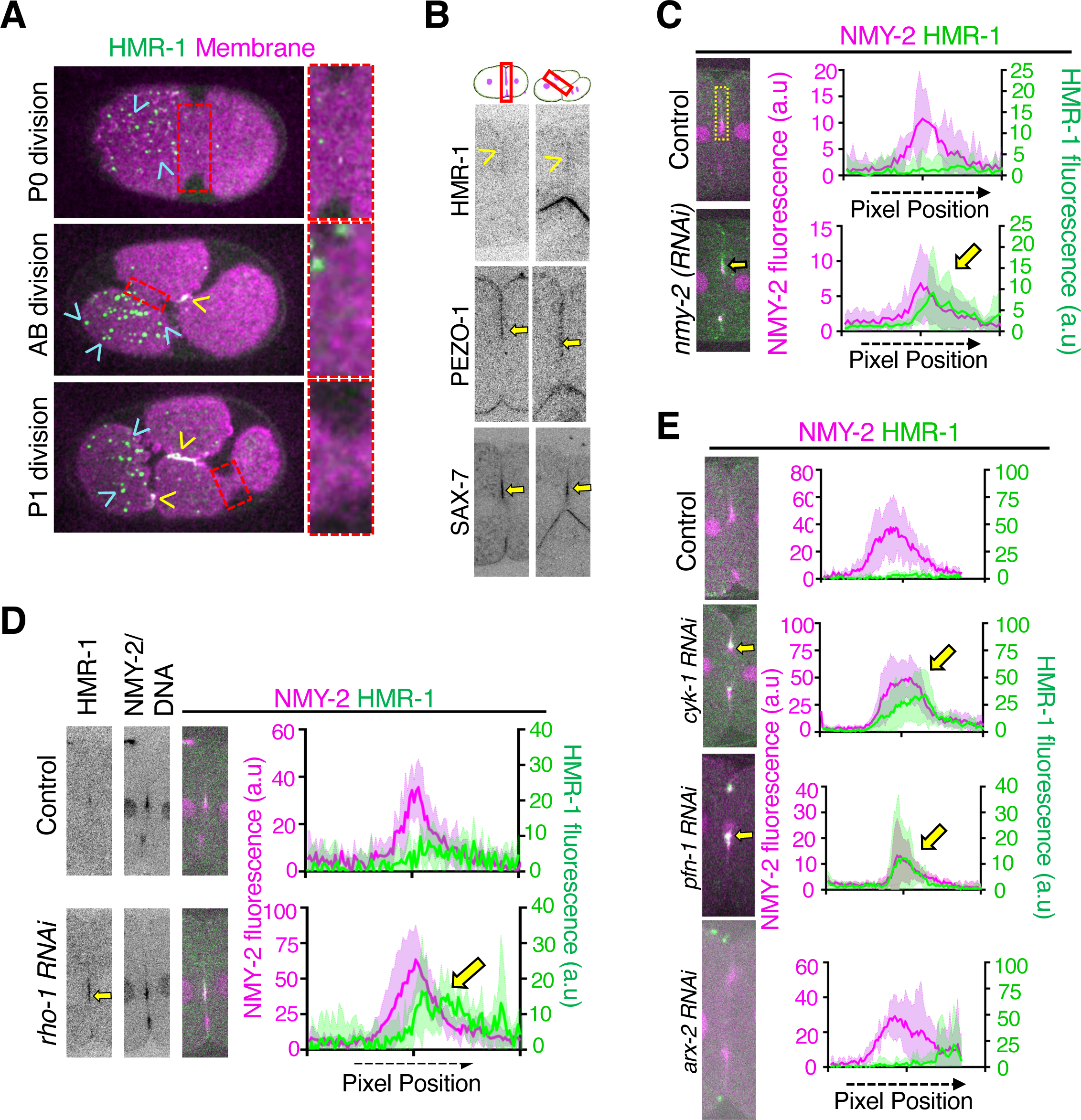
Surface localized HMR-1 is excluded from the contractile furrow zone during cytokinesis. **A.** Cell surface view of embryos co-expressing HMR-1::GFP (green) and mCherry:: PLC1δ-PH (magenta) during P0, AB and P1 cytokinesis. HMR-1::GFP clusters are detected at non-junctional surfaces (cyan arrowhead) and at inter-cellular junctions (yellow arrowhead). The enlarged region of interest (red ROI) indicates the furrow zone undergoing ingression and is devoid of HMR-1::GFP clusters. **B.** Cross-sectional view of furrow zone during cytokinesis in P0 and AB cells expressing HMR-1::GFP, PEZO-1:: mScarlet or SAX-7::eGFP. PEZO-1::mScarlet and SAX-7::eGFP localizes to the ingressing furrow during both P0 and AB-cell division (yellow arrow), but not HMR-1::GFP (yellow arrowhead). **C.** Medial plane of cytokinetic furrows during P0 division in control and *nmy-2(RNAi)* embryos co-expressing NMY-2::mCherry, Histone::mCherry and HMR-1::GFP. Yellow arrows indicate HMR-1::GFP localization in the furrows of *nmy-2(RNAi)* embryos. Plots denote mean ± 95% CI of NMY-2::mCherry and HMR-1::GFP intensities measured in the ingressing furrow as indicated by the yellow ROI in control (n=3) and *nmy-2 (RNAi)* embryos (n=5). **D.** Increased HMR-1::GFP localization in the furrows during P0 division in embryos depleted of RHO-1 (yellow arrows). Plots indicate Mean ±95% CI of NMY-2::mCherry, Histone::mCherry and HMR-1::GFP intensities in the ingressing furrow of control (n=6) and *rho-1* (RNAi) embryos (n=8). **E.** Depletion of CYK-1 but not ARP2/3 complex results in HMR-1 accumulation in the furrow. Equatorial images of furrow ingression during P0 division in embryos co-expressing HMR-1::GFP, Histone::mCherry, and NMY-2::mCherry and depleted of F-actin nucleators as indicated. Mean ±95% CI of NMY-2::mCherry and HMR-1::GFP intensity in the ingressing furrow of control (n=5) and *cyk-1(RNAi)* (n=9), *pfn-1(RNAi)* (n=5), *arx-2(RNAi)* (n=4) embryos are plotted.

### HMR-1 exclusion from the cytokinetic furrow zone is NMY-2 dependent but independent of endocytic trafficking, protein degradation and polarity factors

Earlier investigations demonstrated that the depletion of HMR-1 led to increased NMY-2 dependent contractility at the non-junctional surfaces of the *C. elegans* zygote and early embryos suggesting regulatory role of HMR-1 receptors in inhibiting actomyosin contractility (Padmanabhan et al., 2017b). During cytokinesis, the medial cortex and the ingressing furrow are enriched in NMY-2. Since we found these regions to be devoid of HMR-1, we investigated the possibility of a reciprocal inhibitory mechanism, wherein cortical NMY-2 might also hinder the localization and/or dynamics of HMR-1. We therefore quantified the levels of HMR-1 in embryos partially depleted for NMY-2 (to allow furrow formation and ingression) and co-expressing HMR-1::GFP, NMY-2::mCherry and mCherry:Histone. While we did not detect any HMR-1::GFP fluorescence along the furrow in control embryos, a ∼50% depletion of NMY-2 led to a fivefold accumulation of HMR-1 (Fig 1C; yellow arrow in lower panel) in the cytokinetic furrow.

In addition to sorting of adhesion molecules as cargo, vesicle trafficking during cytokinesis supplements increase in surface area accompanying furrow ingression. (Ai and Skop, 2009; Kakar-Bhanot et al., 2019; Lock and Stow, 2005; Sakaguchi et al., 2015; Skop et al., 2001; Yabuno et al., 2019). This prompted us to investigate the potential role of endocytic trafficking in HMR-1 transport during cytokinesis, disruption of which might lead to an accumulation of HMR-1 in the furrow region. However, we did not detect HMR-1 in the furrow upon RNAi mediated depletion of candidates functioning in the clathrin (*chc-1, unc-11*) or caveolin (*cav-1*) mediated endocytosis or vesicle trafficking (*dhc-1, rab-11.1*) (Fig S1C). CDC-42/PAR mediated A-P polarity maintenance drives the asymmetric localization of HMR-1 as well as asymmetric cell division in *C. elegans* zygote (Cheeks et al., 2004; Hung T.J. and Kemphues K. J., 1999; Motegi and Sugimoto, 2006). To check if this asymmetric HMR-1 distribution influenced its dynamics during cytokinesis, we disrupted A-P polarization pathway (*cdc-42, par-5, par-6*) and found no effect on HMR-1 localization (Fig S1C). Ubiquitination of E-cadherin receptors has also been implicated in E-cadherin trafficking (Niño et al., 2019). We therefore investigated the possibility of HMR-1 turnover during recycling and/or degradation as a mechanism to deplete HMR-1 levels in the cytokinetic furrow. Depletion of the *C. elegans* ortholog of E2 Ubiquitin conjugating enzyme UEV-1 did not result in any observable accumulation of HMR-1 in the cytokinetic furrow (Fig S1C).

Taken together, these results indicate that endocytic trafficking, polarity establishment and ubiquitin mediated proteasomal degradation have negligible role in the cytokinetic furrow being devoid of HMR-1.

### Loss of Formin/CYK-1 allows accumulation of HMR-1 in the cytokinetic furrow

RHO-1, the ortholog of RhoA GTPase in *C. elegans,* activates NMY-2 activity at the furrow zone via phosphorylation of the myosin light chain MLC-4 (Basant and Glotzer, 2018; Matsumura, 2005; Wissmann et al., 1997). While a complete loss of RHO-1 activity severely affected the germline development and prevented any form of furrow ingression, a partial depletion in *rho-1(RNAi)* embryos, allowed formation and constriction of the cytokinetic furrow, albeit at a slower rate. To test if a reduction in RHO-1 would phenocopy *nmy-2(RNAi)* in terms of HMR-1 localization, we partially depleted RHO-1 in embryos expressing HMR-1::GFP and NMY-2::mCherry. We detected a ∼3-fold increase in the levels of HMR-1 at the edge of the furrow in *rho-1(RNAi)* embryos compared to the control (Fig1D, yellow arrow). The cortical F-actin in the *C. elegans* embryonic cell cortex primarily consists of unbranched F-actin filaments polymerized by the diaphanous formin ortholog, CYK-1 and branched actin that appear as punctate structures, nucleated via the ARP-2/3 complex. RHO-1 activates CYK-1 at the division furrow to polymerize F-actin filaments to be assembled into the cytokinetic ring (Otomo et al., 2005; Swan et al., 1998). Additionally, the spatiotemporal regulation of cortical F-actin architecture is orchestrated by various actin binding proteins (ABDs) comprising of crosslinkers, bundling proteins, nucleation promotion factors, severing proteins etc. HMR-1 receptors associate with cortical F-actin via their c-terminal cytoplasmic domain, mediated by adaptor molecules – HMP-2 and HMP-1 (Adams et al., 1998; Costa et al., 1998). We investigated if cortical association with F-actin could restrict HMR-1 mobility into the furrow during cytokinesis. To this end, we disrupted different aspects of cortical F-actin organization by depleting F-actin regulators in embryos co-expressing HMR-1::GFP and NMY-2::mCherry. While HMR-1::GFP signals were not detected in furrows of control embryos, a 10-fold increase in HMR-1 was observed in *cyk-1(RNAi)* embryos or by upshifting temperature sensitive *cyk-1(ts*) mutant embryos to restrictive temperature (Fig 1E-second row, S1D; yellow arrow). CYK-1 requires profilin/PFN-1 bound G-actin to elongate the barbed end of F-actin filaments (Severson et al., 2002). Like in *cyk-1(RNAi)*, *pfn-1(RNAi)* embryos displayed enrichment of HMR-1 in the cytokinetic furrow (Fig 1E-third row, yellow arrow). Since PFN-1 is also required for assembly of branched F-actin, we checked if there was any the role of ARP-2/3 pathway in HMR-1 dynamics during cell division. In contrast to loss of CYK-1, depletion of F-actin regulators that promote branched F-actin architecture (*arx-2, wip-1, wve-1, rac-1*), had negligible effect on the levels of HMR-1 in the furrow during cytokinesis (Fig 1E-last row, S1C). Additionally, we did not detect any effect on HMR-1 localization in ingressing furrows of zygotes depleted of the scaffolding protein, anillin (ANI-1), septin (UNC-59) or cofilin (UNC-60) (Fig. S1C). These results demonstrate that absence of HMR-1 localization at the cytokinetic furrow is mediated by CYK-1.

### Lateral diffusion of HMR-1 clusters during furrow ingression is inhibited by CYK-1

To check if depletion of CYK-1 regulates lateral diffusion of HMR-1 into the newly formed membrane surface during cytokinesis, we analyzed the movement of surface localized HMR-1::GFP clusters as the furrow began to ingress. The point at which surface deformed inwards was taken as ‘time=0’. As described in earlier studies, in control embryos expressing HMR-1::GFP, cortical rotation concomitant with the initiation of furrow ingression dragged HMR-1 clusters along the direction parallel (i.e y-axis) to the contractile furrow zone (y-axis), with a mean speed of 0.31±0.15 μm/s (Fig 2A(a_y_) and 2B, blue arrow in brown ROI). Following this initial movement along the y-axis, HMR-1 clusters paused and began moving perpendicular, along the x-axis (+100s to +200s) towards the ingressing furrow, with at a mean speed of 0.13±0.06 μm/s (Fig. 2A(a_x_), 2B, blue arrow in orange ROI and 2C). Upon a partial depletion of CYK-1 we observed ∼50% reduction in speeds of HMR-1 clusters (0.18±0. 07 μm/s) along the Y-axis (Fig 2A(b_y_) and 2B, red arrow in brown ROI). Subsequent movement of these HMR-1 clusters along x-axis into the furrow however, displayed a slight but significant increase in its speed 0.15±0.05 μm/s (Fig 2A(b_x_), 2B red arrow in orange ROI and 2C). We then measured the mean square displacement (MSD) of HMR-1 clusters during the period (+100s - +200s) (Fig. 2D). Our estimated diffusion coefficient value of 0.013±0.001 μm^2^ s^-1^ for HMR-1 clusters in control embryos are in agreement with values reported for E-cadherin in other systems (Adams et al., 1998; Biswas et al., 2015; Kusumi et al., 1993). Depletion of CYK-1 led to a ∼20 fold increase in the diffusion coefficient value (0.19±0.13 μm^2^ s^-1^) (Fig. 2E). A value greater than 1 (∼1.59±0.22) for the diffusion exponent ‘α’ for HMR-1 clusters in control embryos suggested super-diffusion. In *cyk-1(RNAi)* embryos, however, HMR-1 clusters exhibited ‘sub-diffusion’ (‘α = ∼0.76±0.20) as they moved into the cytokinetic furrow zone (Fig 2F). Taken together our results show that CYK-1 inhibits lateral movement of HMR-1 clusters into the furrow zone as well as the newly formed membrane surface following invagination.

**Figure 2:**
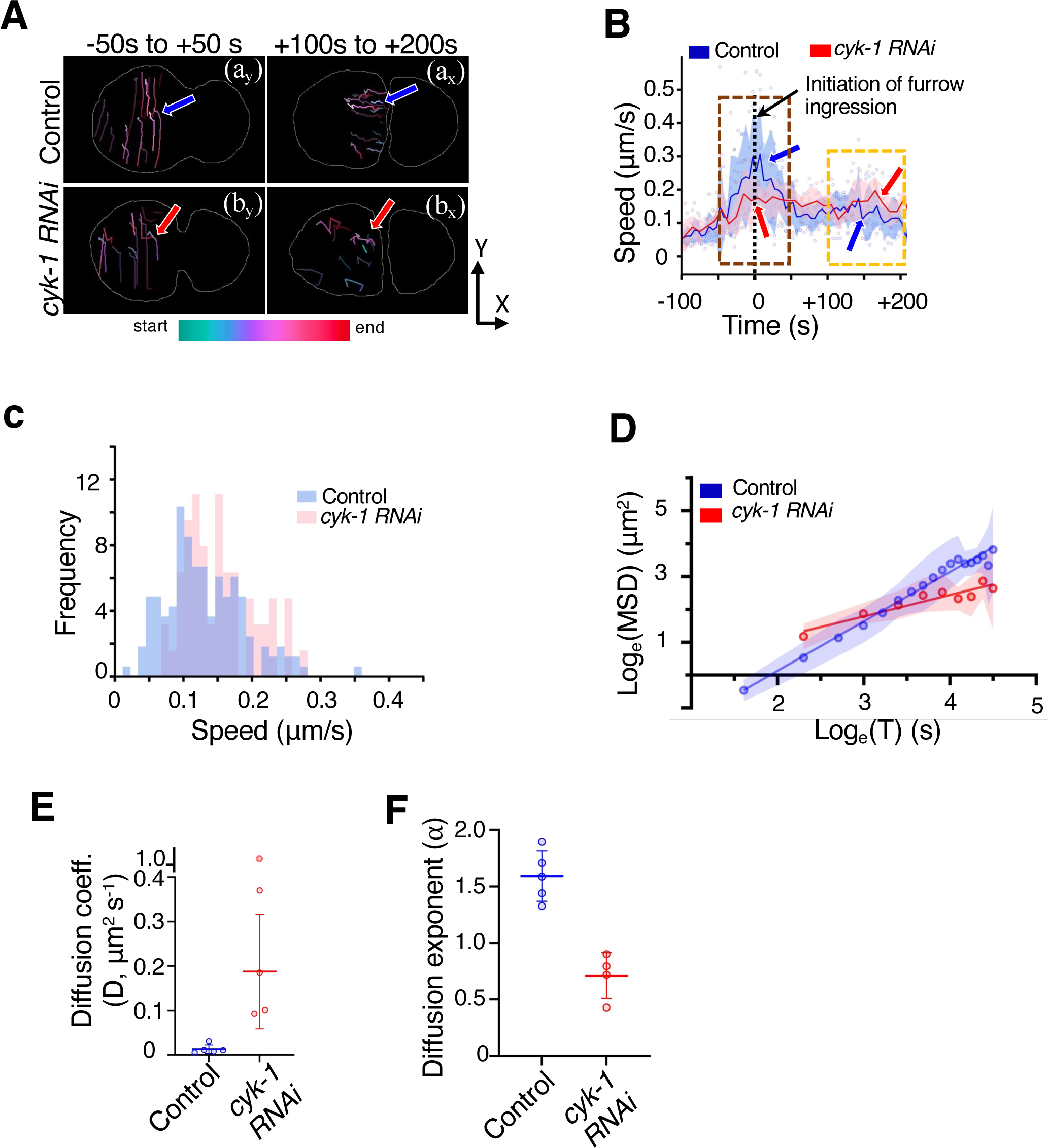
CYK-1 depletion enhances the lateral movement of HMR-1 clusters during cytokinetic furrow ingression. **A.** Depletion of CYK-1 attenuates the initial cortical movement of HMR-1 clusters parallel to furrow (-50s to +50s), but not the subsequent perpendicular migration into the furrow(+100s to +200s). Color coded tracks indicate the direction of movement of few selected HMR-1::GFP clusters during control and *cyk-1(RNAi)* cytokinesis. **B.** Plot depicting instantaneous speed of HMR-1::GFP clusters on the cell surface (vs) the time during furrow ingression in control (blue) and *cyk-1(RNAi)* (red) zygotes. CYK-1 depletion slows down the early cortical movement of HMR-1 clusters (brown ROI at -50s to +50s, red arrow), but significantly increase HMR-1 speeds during later time (orange ROI at +100s to +200s, red arrow). **C.** Normalized histogram of HMR-1::GFP cluster speeds during cortical flow of puncta in the furrow cleavage region (time: ∼+80s to +200s) for control (blue) and *cyk-1(RNAi)* (red) embryos (n=5). **D.** The linear fit of the log_e_(MSD) vs log_e_(time) for the mean of HMR-1::GFP cluster movement during +100s to +200s in control (blue) and *cyk-1*(*RNAi)* (red). **E.** and **F.** The diffusion coefficient (D) and diffusion exponent (⍺) of HMR-1 clusters for individual embryos analysed in (Fig 2D).

### Cortical organization of CYK-1 polymerized F-actin filaments restrict the movement of HMR-1 clusters at the cytokinetic furrow zone

During cytokinesis, various acting binding and regulatory proteins re-organize, crosslink and bundle unbranched F-actin filaments at the medial cortex into a parallel architecture (Sobral et al., 2021). Given that surface localized clusters of HMR-1 associate with F-actin during cortical actomyosin flows during polarization and cytokinesis (Padmanabhan et al., 2017b), we investigated if this association between HMR-1 and F-actin filaments might regulate HMR-1 dynamics at the division furrow. To this end, we imaged the surface of embryos co-expressing HMR-1::GFP and the F-actin marker Lifeact fused to RFP (Lifeact::RFP). F-actin filaments get organized in a parallel arrangement in control embryos (Fig 3A, red ROI) (Leite et al., 2020; Li and Munro, 2021; Reymann et al., 2016). In these embryos HMR-1 clusters were detected only along the outer edge of the region marked by the parallel F-actin filaments with less than 2 clusters in the furrow zone (Fig 3A i(a) and 3B). Depletion of CYK-1 resulted in significant reduction in the density of unbranched F-actin filaments while punctate structures of ARP-2/3 nucleated branched F-actin structures were visible on the cortical surface (Fig 3A, *cyk-1(RNAi)).* Compared to control embryos, we detected significant increase in the number of HMR-1 clusters in the furrow region in *cyk-1(RNAi)* embryos (12.2±3.8 clusters in *cyk-1(RNAi)*) (Fig. 3A ii(a), 3B). Depletion of ARX-2 abolished ARP2/3 dependent branched F-actin structures which led to an increased density of unbranched F-actin filaments. In *arx-2(RNAi)*, HMR-1 clusters were found to be displaced towards the anterior end and not more than 1-2 clusters were detected in the furrow zone during division (Fig. 3A iii(a), 3B). The accumulation of HMR-1 clusters upon profilin depletion (18±5.8 clusters in *pfn-1(RNAi)*) indicated that the spatial inhibition of HMR-1 localization at the furrow zone was mediated by the F-actin polymerization activity of CYK-1 (Fig 1E, 3A and 3B).

**Figure 3:**
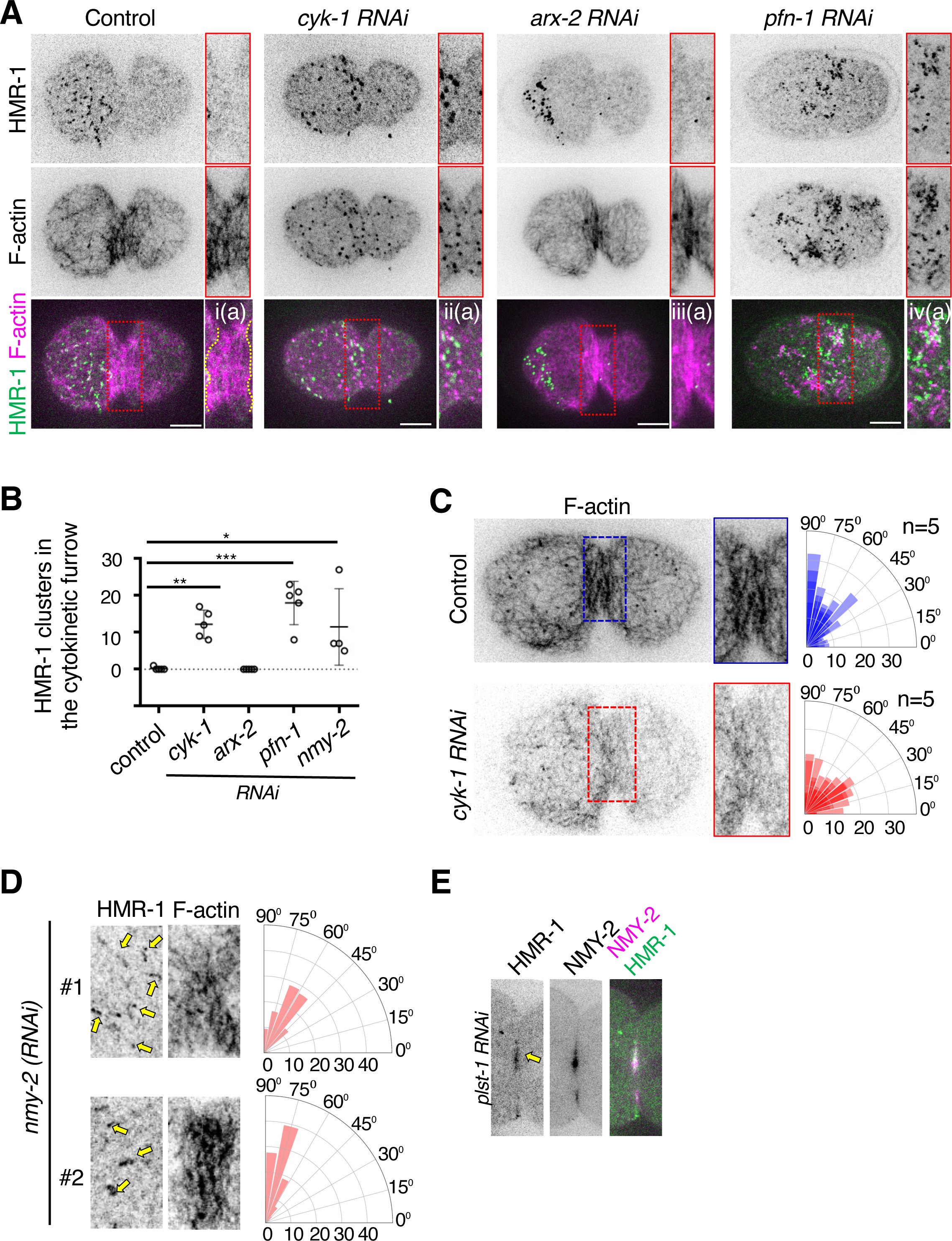
Parallel alignment of F-actin structures at the furrow compact to restrict HMR-1 mobility during cytokinesis associated cortical flows. **A.** Surface view of first embryonic division in control, *cyk-1(RNAi), arx-2(RNAi),* and *pfn-1(RNAi)* zygotes co-expressing HMR-1::GFP and lifeACT::RFP. Enlarged furrow region (red ROI) shows the presence of HMR-1::GFP accumulation in *cyk-1(RNAi), pfn-1(RNAi)* furrow zone but not in that of control and *arx-2 (RNAi)* embryos. **B.** Quantification of the number of HMR-1 clusters in the cytokinetic furrow zone of embryos listed in Fig 3A and 3D (n=5). Error bars are mean ± 95% CI (*p < 0.05, **p < 0.005, ***p < 0.0001, ANOVA test) **C. and D.** Cortical F-actin filaments in control embryos display compact alignment along the ingressing furrow oriented perpendicular to the embryonic axis (blue ROI in Fig 3C top panel). F-actin filaments in *cyk-1(RNAi)* and *nmy-2(RNAi)* embryos are randomly oriented (red ROI in Fig 3C lower panel in 3D). Radial histogram shows the relative normalized filament orientation of indicated embryos. (n=5). **D.** The number of HMR-1 clusters accumulated in the furrow is directly proportional to the disruption of F-actin ortienation (Fig 3D yellow arrows in #1 top panel) **E.** PLST-1 depletion leads to HMR-1 accumulation in the furrow zone (yellow arrow). Equatorial image of furrow ingression in a *plst-1(RNAi)* zygote co-expressing HMR-1::GFP and NMY-2::mCherry.

Multiple mechanisms have been proposed to function in orienting F-actin filaments into a compact and parallel architecture with majority of the filaments oriented 90⁰ to the embryonic axis (Henson et al., 2017; Maupin and Pollard, 1986; Leite et al., 2020). To better understand how local F-actin organization might dictate HMR-1 mobility, we analyzed the orientation of F-actin filaments in the furrow zone in control and *cyk-1(RNAi)* embryos. In agreement with earlier studies, F-actin filaments get aligned and bundled into a parallel architecture that was perpendicular to the embryonic axis (Fig. 3C, blue ROI). In contrast, fewer F-actin filaments were observed in *cyk-1 (RNAi)* embryos and those filaments were disordered in their orientation (Fig. 3C, red ROI). NMY-2 and the F-actin crosslinker, plastin/PLST-1 align and compact cortical F-actin during cytokinesis (Leite et al., 2020). *nmy-2(RNAi)* embryos displayed a modest accumulation of HMR-1 in the furrow (Fig 1C). We therefore inquired if the F-actin architecture in the furrow zone had any bearing on the extent of HMR-1 accumulation. In dividing embryos depleted of NMY-2 and expressing HMR-1::GFP and Lifeact::RFP, we observed (as previously reported) a disruption of F-actin organization in the furrow zone (Fig. 3D). On an average we detected ∼11.5±5.2 clusters upon NMY-2 depletion (Fig 3B). We also noticed that the extent of F-actin orientation depended on the penetrance of *nmy-2(RNAi)*, with higher knockdown of NMY-2 displaying highly disordered F-actin filaments. Additionally, there seemed to be a direct correlation between the extent of disruption in F-actin organization and the medial accumulation of HMR-1, with higher number of HMR-1::GFP puncta in the furrow zone of embryos showing highly disordered F-actin filaments. (Fig 3D, panel #1, yellow arrows)

Plastin (PLST-1) has been shown to crosslink linear-F-actin filaments and plays a key function in cortical actomyosin connectivity during polarization and cytokinesis (Ding et al., 2017)., Embryos lacking PLST-1, exhibited defects in F-actin bundling and delayed cytokinetic furrow ingression, phenotypes that are attributed to defective crosslinking of F-actin filaments at the furrow zone (Sobral et al., 2021). Similar to *cyk-1 (RNAi)* and *nmy-2(RNAi)*, we detected HMR-1::GFP in the ingressing furrows of *plst-1(RNAi)* embryos (Fig. 3E, yellow arrow).

These observations indicate a mechanism where the compact architecture of parallel F-actin filaments corralled HMR-1 from moving along with the cortical flow into the furrow zone. Perturbation of this F-actin filament organization at the medial cortex allows HMR-1 to diffuse into the ingressing furrow.

### HMR-1 associates with cortical F-actin via catenins during cytokinesis associated cortical flows

The *C. elegans* orthologs of β- and α- catenin - HMP-2 and HMP-1 respectively, mechanically link the C-terminal cytoplasmic domain of HMR-1 to cortical F-actin at cell-cell junctions (Choi et al., 2015; Costa et al., 1998; Kang et al., 2017). Therefore, perturbing function of HMP-2 /HMP-1 decouples F-actin filaments from HMR-1 receptors, resulting in disruption of junctional integrity during epithelial morphogenesis (Choi et al., 2015; Costa et al., 1998; Kang et al., 2017; Loveless and Hardin, 2012). Biochemical characterization of HMP-1 indicated its constitutive association with HMP-2 and F-actin (Kang et al., 2017). Given that, non-junctional surface localized HMR-1 in the early *C. elegans* embryos associate with cortical F-actin (Padmanabhan et al., 2017b) during cortical flows, we investigated if this association was mediated through HMP-2 /HMP-1, similar to junctional HMR-1. In embryos co-expressing Lifeact::RFP and HMP-2::GFP or HMP-1::GFP (at endogenous locus), we observed HMP-2 and HMP-1 localize to cell-cell junctions between AB and P1 cells (yellow arrows in Fig S4A and S4B) as well as clusters at non-junctional surfaces (yellow arrows in Fig 4A and 4B red ROI). The surface localization of HMP-2 and HMP-1 mirrored that of non-junctional HMR-1 seen at the cell surface (Padmanabhan et al., 2017b). Co-localization analysis revealed that as previously reported for HMR-1, non-junctional HMP-2::GFP and HMP-1::GFP clusters co-localized with F-actin filaments during initiation cortical flows (yellow arrows in enlarged ROIs in Fig 4A, 4B). To test if the localization of HMP-2 and HMP-1 at the non-junctional cell surface is contingent upon its association with the cytoplasmic domain of HMR-1, we depleted HMR-1 in these embryos. Loss of HMR-1 completely abrogated HMP-2 and HMP-1 from both cell-cell junctions and the non-junctional surfaces (Lower panels in Fig 4A and 4B, S4A and S4B). Taken together we find that catenins- HMP-2 and HMP-1 are recruited to the cell surface by HMR-1where they mediate the mechanical association of HMR-1 with F-actin.

**Figure 4.**
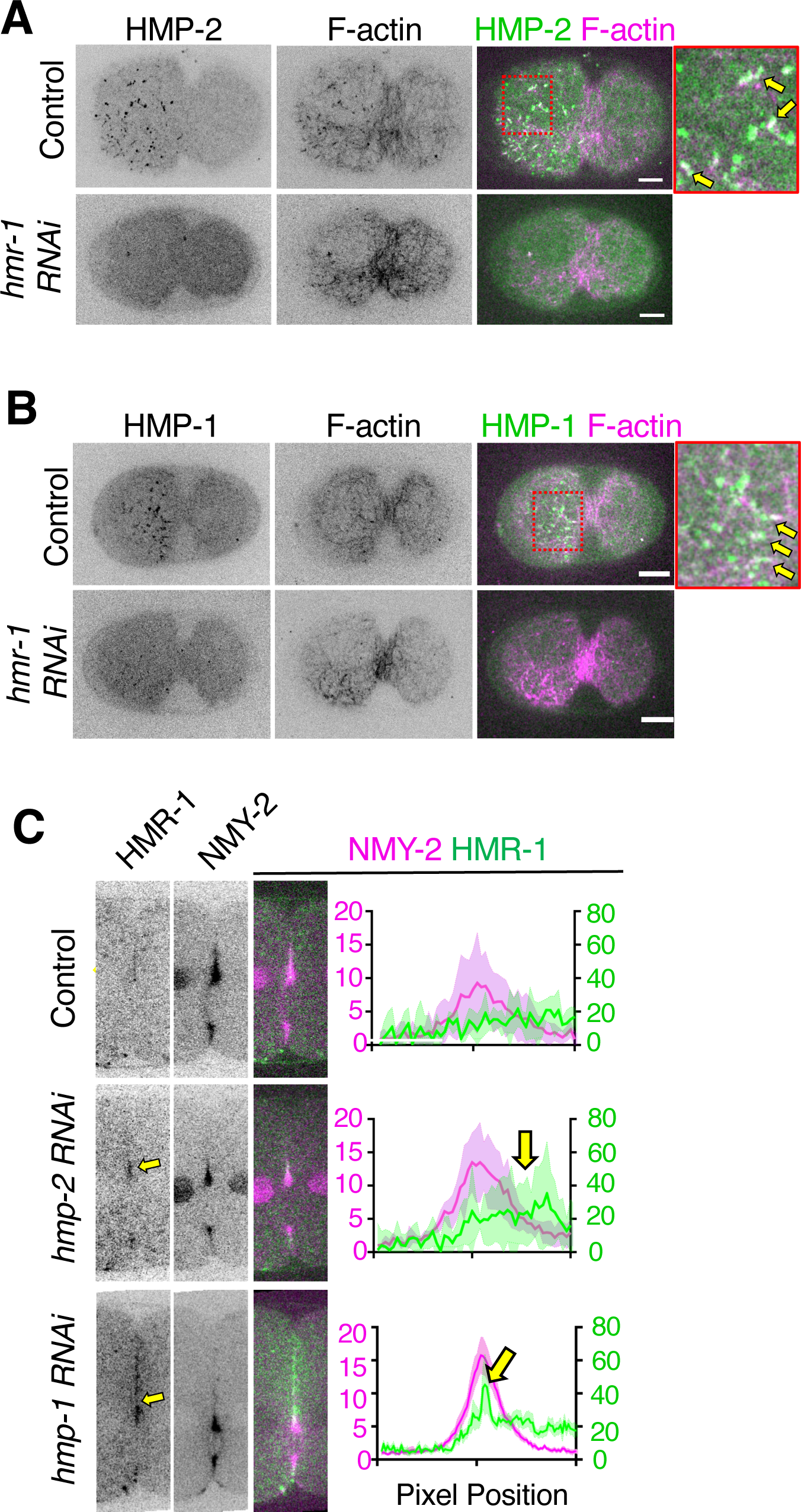
– Spatial exclusion of non-junctional HMR-1 during cytokinesis is regulated by HMP-2/HMP-1 mediated association with cortical F-actin. **A. and B.** Surface view of one-cell stage embryos undergoing cytokinesis and co-expressing HMP-2::GFP or HMP-1::GFP and lifeACT::RFP. HMP-2::GFP and HMP-1::GFP clusters at the cell surface colocalises with F-actin in control embryos (yellow arrows in red ROI). Surface localization of HMP-2/HMP-1 is disrupted in *hmr-1(RNAi)* embryos. **B.** Depletion of HMP-2 or HMP-1 result in the accumulation of HMR-1::GFP in the ingressing furrow. Medial confocal plane of cytokinetic furrows from P0 division in control, *hmp-2(RNAi)* and *hmp-1(RNAi)* embryos co-expressing NMY-2::mCherry, Histone::mCherry and HMR-1::GFP. Plot shows mean ± 95%CI of NMY-2::mCherry and HMR-1::GFP intensity in the ingressing furrow in control (n=4) *hmp-2(RNAi)* (n=5) and *hmp-1(RNAi)* (n=11) embryos. Yellow arrows indicate HMR-1::GFP localization in the ingressing furrow.

### HMP-2/HMP-1 mediated engagement with F-actin restricts HMR-1 localization at the division furrow

Mechanical linkage with F-actin has been shown to regulate E-cadherin mobility on the cell surface (Chandran et al., 2021; Sako et al., 1998; Wu et al., 2015). Therefore, we examined the possibility that HMP-2/HMP-1 mediated linkage between HMR-1 to F-actin might be restricting its dynamics at the furrow zone during cytokinesis. We measured the levels of HMR-1 in the furrow region of embryos expressing HMR-1::GFP and NMY-2::mCherry, upon depletion of HMP-2 or HMP-1. Compared to control embryos, ∼4 fold increase in HMR-1 levels were observed in the furrows of *hmp-1(RNAi)* and *hmp-2 (RNAi)* embryos (Fig. 4C, yellow arrows). These results demonstrate that HMP-2 and HMP-1 connect HMR-1 to the cortical F-actin and this linkage is essential to prevent HMR-1 from moving into the furrow zone during cytokinesis associated cortical flow.

## DISCUSSION

The physical connection between HMR-1 and F-actin leads to HMR-1 being carried along during cortical actomyosin flows. The localization of HMR-1 on the cell surface mirrors the direction of the cortical flow, as evidenced by the accumulation of HMR-1 in the anterior half of the *C. elegans* zygote after polarization. Cortical F-actin flows towards the cytokinetic furrow have been observed in various cell types, including the *C. elegans* embryos, suggesting that this flow should result in the localization of HMR-1 in the furrow zone (Cao and Wang, 1990; Pinheiro et al., 2017; Reymann et al., 2016). Contrary to this expectation, during the initial embryonic divisions in *C. elegans*, the flow of F-actin into the cytokinetic furrow does not bring in HMR-1 (Padmanabhan et al., 2017b). How is HMR-1 restricted from furrow zone during cytokinesis?

In this study we show that the spatial inhibition of HMR-1 clusters at the cytokinetic furrow is mediated by, (1) parallel architecture of CYK-1 polymerized unbranched F-actin filaments at the medial cortex, (2) catenin-mediated physical linkage between the HMR-1 and F-actin. During cytokinesis, cortical F-actin polymerized as unbranched filaments gets compacted into an parallel architecture. These findings support the hypothesis that such an F-actin organization associates with the cytoplasmic domain of HMR-1. This impedes HMR-1 mobility and prevents its localization to furrow zone. In summary, our studies provide a plausible explanation for the mutual antagonism between E-cadherin mediated cell adhesion and the cortical actomyosin based cell division machineries.

## ACKNOWLEDGEMENTS

This work was supported by DBT-Wellcome India Alliance Fellowship (**IA/I/18/1/503624)** to A.P, and core funding support from the Trivedi School of Biosciences, Ashoka University. We acknowledge the infrastructure support from the Microscopy Core facility at Ashoka University. We thank Andy Golden and Ronen Zaidel-Bar for sharing strains. Some strains were provided by the CGC, which is funded by NIH Office of Research Infrastructure Programs (P40 OD010440).

## AUTHOR CONTRIBUTIONS

A.P conceived the project. D.M, M.R and A.P designed and performed the experiments. D.M and M.R analysed the data with A.P’s input. A.P wrote the manuscript with input from D.M and M.R.

---

**Figure S1.**
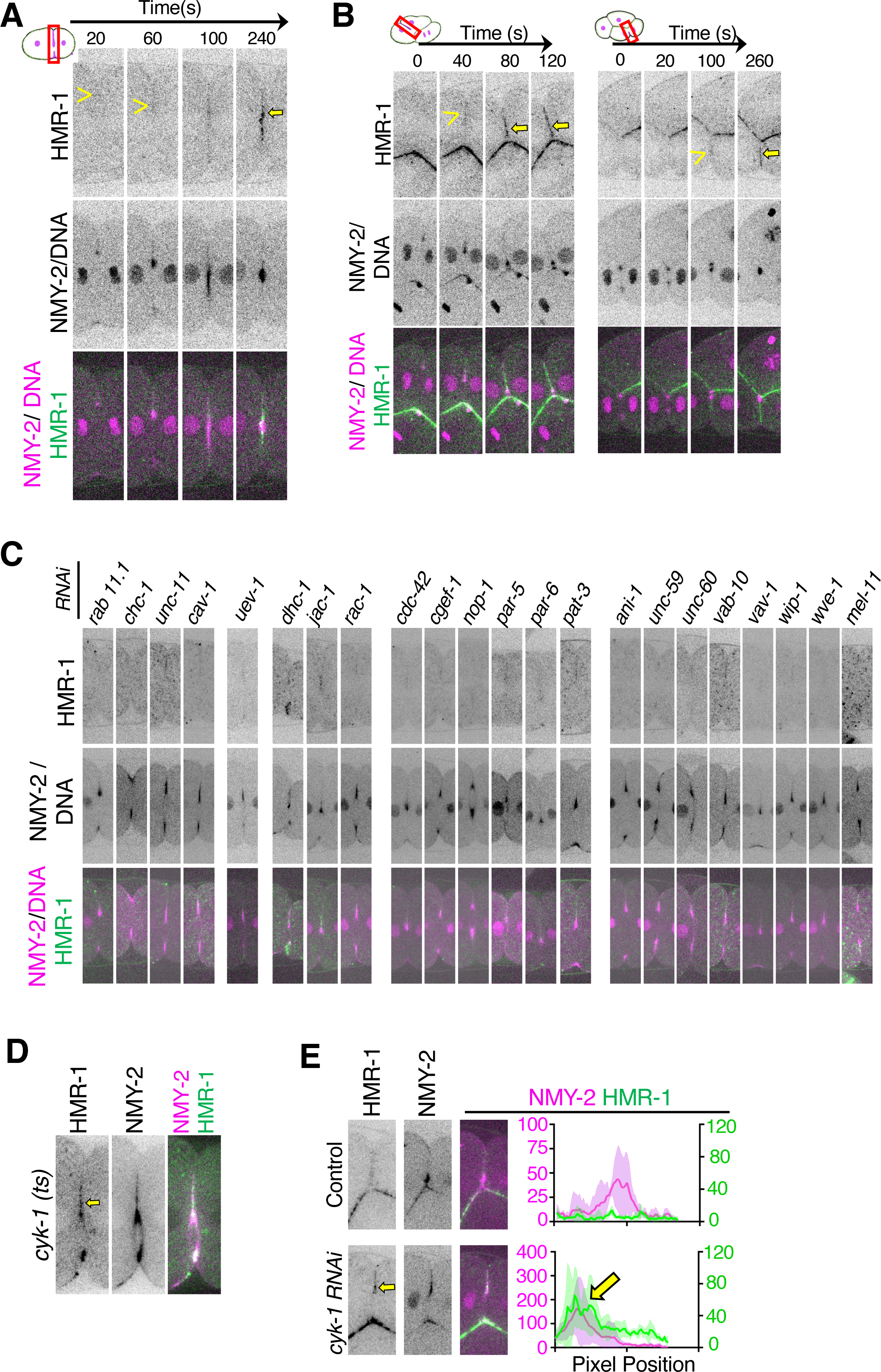
**A.** Time-lapse images of furrow ingression during the first division (P0 cell) in an embryo co-expressing NMY-2::mCherry Histone::mCherry and HMR-1::GFP. The yellow arrow head shows the absence of HMR-1::GFP in the ingressing furrow but following cytokinesis completion, HMR-1::GFP localizes to the inter-cellular boundary (yellow arrow). **B.** Time-lapse images of furrow ingression during the AB cell division (left panel) and P1 division (right panel) in an embryo co-expressing NMY-2::mCherry, Histone::mCherry and HMR-1::GFP. Similar to P0 cytokinesis, HMR-1::GFP is absent in the ingressing furrow (yellow arrowhead) but localizes to the inter-cellular boundary upon completion of cytokinesis (yellow arrow). **C.** Medial region of the cytokinetic furrow during P0 division in embryo co-expressing NMY-2::mCherry, Histone::mCherry and HMR-1::GFP. Candidate RNAi screening of the selected genes are indicated. **D.** Equatorial image of furrow ingression during the P0 cytokinesis of *cyk-1(ts)* embryo co-expressing HMR-1::GFP and NMY-2::mCherry. Acquired at the restrictive temperature (25°C), HMR-1::GFP is localized in the furrow upon loss of CYK-1 activity (yellow arrow). **E.** Equatorial image of furrow ingression during the AB-cell cytokinesis, co-expressing NMY-2::mCherry Histone::mCherry and HMR-1::GFP in control (top panel) and *cyk-1(RNAi)* (bottom panel). The plot shows mean ± 95%CI of NMY-2::mCherry and HMR-1::GFP intensities in the ingressing furrows. The yellow arrow indicates accumulation of HMR-1::GFP in the cytokinetic furrow of *cyk-1(RNAi)* embryos. (n=4)

**Figure S4.**
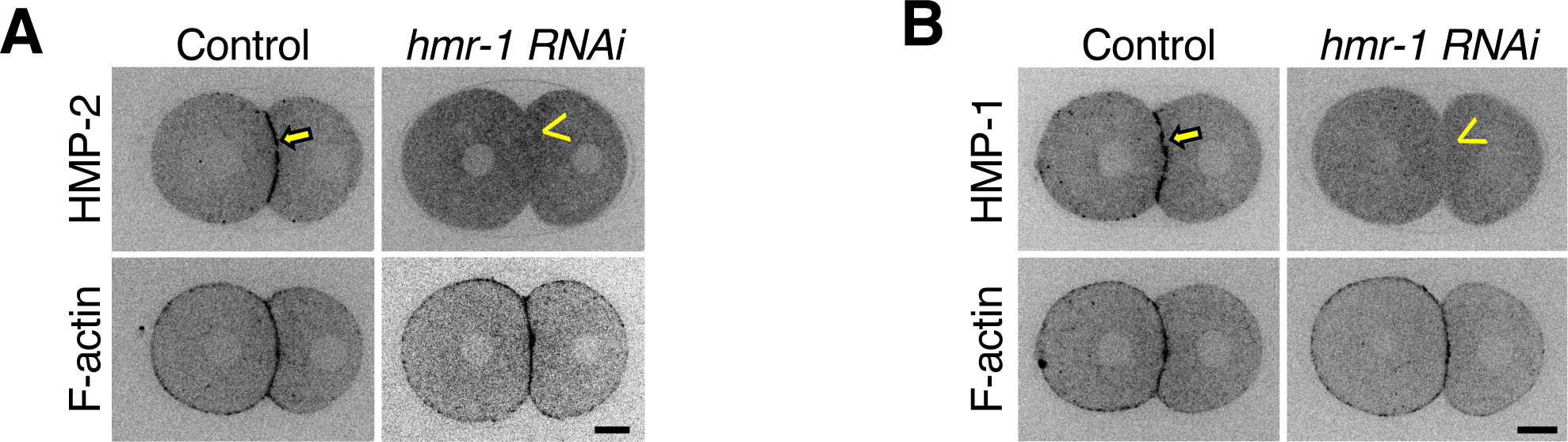
**A. and B.** Equatorial image of furrow ingression in two-cell staged embryo co-expressing HMP-2::GFP or HMP-1::GFP and Lifeact::RFP. The yellow arrow shows HMP-2::GFP or HMP-1::GFP localized at the inter-cellular boundary. Junctional localization of HMP-2::GFP and HMP-1::GFP is abrogated in *hmr-1(RNAi)* embryos (yellow arrowhead).

## MATERIALS AND METHODS

### Strains

Table of *C. elegans* strains used in this study are listed in Table S1. All strains were grown at 20⁰ C on nematode growth media (NGM) seeded with OP50 *E.coli*.

**Table S1.**
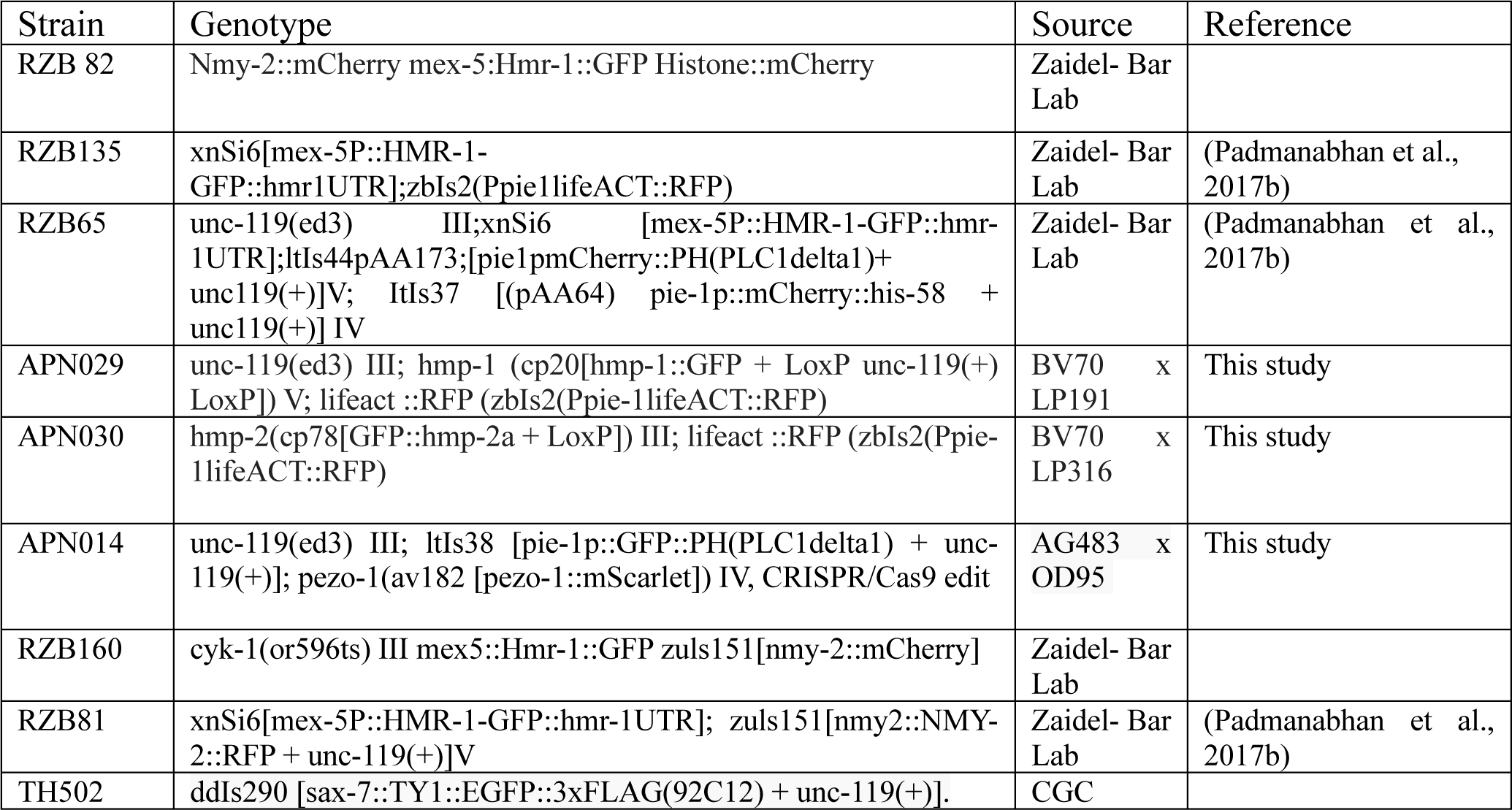

### RNA interference (RNAi)

RNAi was carried out through feeding or dsRNA microinjection. L4-staged hermaphrodites grown in OP50 *E. coli* were collected, washed and transferred to appropriate RNAi plates (Ahringer library) or injected with appropriate dsRNA at a 0.8-1.0 μg/μl.

Details of primers and template used for dsRNA synthesis is indicated in Table S2

**Table S2.**
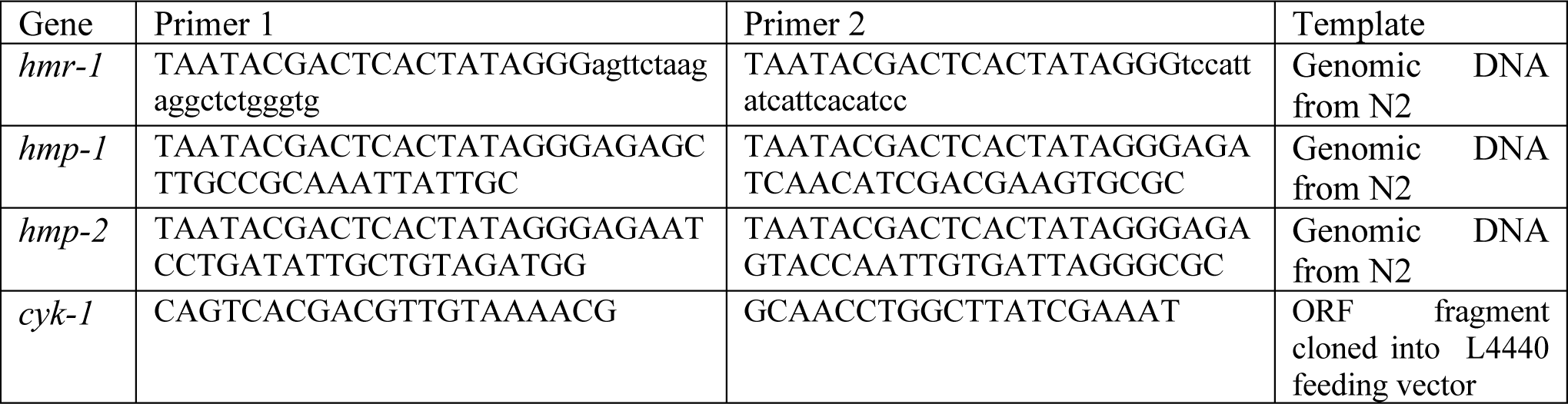

In-vitro synthesized dsRNA for *hmr-1,hmp-1,hmp-2* and *cyk-1* were injected into the L4 stage hermaphrodites using FemtoJet Eppendorf mounted on a IX53 Olympus microscope. 36-48 hours following RNAi treatment, individual hermaphrodites were dissected, and embryos at zygotic stage were collected, mounted on 3% agarose pads, and imaged. RNAi knockdown of *cyk-1, nmy-2* and *plst-1* were confirmed by delayed closure of cytokinetic furrow. *hmr-1, hmp-1, and hmp-2* knockdowns were validated by observing lethality and morphogenesis defects in the dissected embryos.

### Live imaging and Image Acquisition

All imaging was carried out at 20-21⁰C on an Olympus IX83 Inverted Microscope equipped with a 100x UPlanSApo 1.40 NA objective equipped with CSU-W1 spinning-disk confocal head (Yokogawa Corporation, Tokyo, Japan). Embryo samples were excited at 488 and 561 nm using OBIS Coherent laser system and images were acquired using a PrimeBSI camera (2×2 binning), or emCCD Photometric evolve delta camera (BIN1). Image acquisition was controlled by Olympus CellSens Dimension software.

### Image Analysis

Image analysis and quantifications were carried out using Fiji, Python and MS-Excel. Prism 6 software (GraphPad) was used for graph plotting and statistical analysis.

### Intensity line scan plot analysis

To estimate the intensity profile of NMY-2::mCherry and HMR-1::GFP in the ingressing furrow, 5 pixels wide and 100- or 50-pixels long ROI was drawn along the ingressing furrow (yellow ROI, Fig. 1C). Along the ROI, fluorescence intensities of both channels were measured. Cytosolic background in both channels, measured from a region away from the ingressing furrow, were subtracted from furrow intensities. Mean ± 95%CI intensities were plotted.

### Orientation of actin filaments

To analyze F-actin filament orientation, an ROI was selected in the cytokinetic region of surface plane of the embryos expressing Lifeact::RFP. For the ROI, the filament orientation was extracted using the OrientationJ (OJ) Distribution plugin in Fiji(Fonck et al., 2009; Püspöki et al., 2016; Rezakhaniha et al., 2012).

The parameters value used in the OJ plugin are Orientation: degree (-90° to 90°), Gradient: cubic spline, Local window σ=5 pixels (scale:1 pixel=0.16 micron), Minimum energy:0%, Minimum coherency:1%. Custom made Python code was used to extract the absolute value of the filament orientation data (0° to 90°) using the OJ data. The relative Normalized Histogram was calculated for each embryo at -50s and +50s (considered the start of furrow ingression as time ‘0’ for all embryos). Relative normalization was done for the selected ROIs of the embryo at a specific frame. The radial histogram of control and *cyk-1* is a superimposed data for all embryos in respective ROIs of different RNAi experiments (n=5). Radial histogram for *nmy-2 (RNAi*) are plotted for individual embryos.

### Analysis of surface mobility of HMR-1 clusters

To measure the speed of the HMR-1::GFP puncta during cytokinesis, an ROI was selected near the ingressing furrow and the TrackMate plugin in Fiji was used to track the puncta (Ershov et al., 2022). LOG detector was used to select the HMR-1::GFP puncta in the ROI, using the diameter of blob∼0.9 µm and threshold value(∼350 to ∼450) for 16-bit image-type. The LAP tracker was used to track each puncta’s trajectory. The linking distance used for each control embryo (each frame of video:5s) is 5 µm and for *cyk-1(RNAi)* embryos (each frame of video:10s) is 10 µm. The penalties used are Std intensity ch 1:1.0, Median intensity ch 1: 1.0, Visibility: 1.0, Radius: 1.0, Signal/Noise ratio:1. No gap closing distance was used. The tracking data were obtained from the TrackMate as Spots, edge, and Track statistics (.csv) files.

The speed and its x,y-component are taken for control and *cyk-1(RNAi)* using the speed value(extracted from link statistics file generated from TrackMate), the x and y coordinates value of HMR-1::GFP puncta (extracted from spot statistics file generated from TrackMate) that forms a link in two consecutive frames(considered the start of furrow ingression as time ‘0’ for all embryos). For each embryo, the mean speed of all punctae at a given frame is calculated and the mean speed of 5 embryos at each frame is plotted using custom Python code.

To find the diffusion coefficient and diffusion exponent values of puncta during the ingression of Hmr-1::GFP puncta in the furrow ingression region, the puncta were tracked from ∼80s till ∼200s. The Mean square displacement (MSD) for all the tracks of puncta was calculated using custom Python code and the MSD vs track time was plotted. The values of slope and y-intercept of the best fit were used to find the value of diffusion coefficient and diffusion exponent of puncta. The diffusion coefficient and exponent were also calculated from the best fit (Linear regression) for each embryo in control and *cyk-1(RNAi)*.

### Count of HMR-1::GFP clusters in the cytokinetic ring

To estimate the number of HMR-1::GFP clusters at the cytokinetic furrow zone of control, *cyk-1(RNAi), nmy-2(RNAi), pfn-1(RNAi), and arx-2(RNAi)* embryos, in Fiji, the background was subtracted, filtered, and thresholded. After thresholding, images were watershed and the number of HMR-1::GFP clusters were counted using particle analysis plugin.

